# Bispecific antibodies combine breadth, potency, and avidity of parental antibodies to neutralize sarbecoviruses

**DOI:** 10.1101/2022.11.11.516125

**Authors:** Laura Radić, Kwinten Sliepen, Victor Yin, Mitch Brinkkemper, Joan Capella-Pujol, Angela I. Schriek, Jonathan L. Torres, Sandhya Bangaru, Judith A. Burger, Meliawati Poniman, Ilja Bontjer, Joey H. Bouhuijs, David Gideonse, Dirk Eggink, Andrew B. Ward, Albert J. R. Heck, Marit J. Van Gils, Rogier W. Sanders, Janke Schinkel

## Abstract

SARS-CoV-2 mutational variants evade humoral immune responses elicited by vaccines and current monoclonal antibody (mAb) therapies. Novel antibody-based treatments will thus need to exhibit broad neutralization against different variants. Bispecific antibodies (bsAbs) combine the specificities of two distinct antibodies into one antibody taking advantage of the avidity, synergy and cooperativity provided by targeting two different epitopes. Here we used controlled Fab-arm exchange (cFAE), a versatile and straightforward method, to produce bsAbs that neutralize SARS-CoV and SARS-CoV-2 variants, including Omicron and its subvariants, by combining potent SARS-CoV-2-specific neutralizing antibodies with broader but less potent antibodies that also neutralize SARS-CoV. We demonstrate that the parental IgG’s rely on avidity for their neutralizing activity by comparing their potency to bsAbs containing one irrelevant “dead” Fab arm. We used single particle mass photometry to measure formation of antibody:spike complexes, and determined that bsAbs increase binding stoichiometry compared to corresponding cocktails, without a loss of binding affinity. The heterogeneous binding pattern of bsAbs to spike (S), observed by negative-stain electron microscopy and mass photometry provided evidence for both intra- and inter-spike crosslinking. This study highlights the utility of cross-neutralizing antibodies for designing bivalent or multivalent agents to provide a robust activity against circulating variants, as well as future SARS-like coronaviruses.

## Introduction

Severe acute respiratory syndrome coronavirus 2 (SARS-CoV-2) is the cause of the ongoing COVID-19 pandemic, which has up until now resulted in more than 625 million cases and 6.5 million deaths worldwide (https://covid19.who.int/). SARS-CoV-2 has a lower infection fatality rate but is considerably more transmissible than Severe acute respiratory coronavirus (SARS-CoV), which caused a smaller outbreak around 2003 ^1^. Both viruses belong to the *Sarbecovirus* subgenus of the family *Coronaviridae* ^2^. Highly effective SARS-CoV-2 vaccines have been deployed, but there is an inevitable need to develop additional treatment options, especially for certain risk groups, such as immunocompromised adults. Monoclonal antibodies (mAbs) represent some of the most promising candidates for prophylaxis and therapy against viral infections, since their safety and efficacy has been repeatedly shown, e.g. for prevention of RSV infection in premature infants ^3^ and in successfully controlling the Ebola virus epidemic in 2018 ^4,5^.

Numerous potent neutralizing antibodies (NAbs) have been isolated from COVID-19 convalescent patients and extensively characterized ^6–16^. The majority of these NAbs target the receptor binding domain (RBD) of the trimeric spike (S) protein (S-trimer), the main glycoprotein on the viral surface, which both SARS-CoV and SARS-CoV-2 use to bind their host receptor, angiotensin-converting enzyme 2 (ACE2). Antibodies can utilize their two Fab arms to simultaneously bind one or two of the three RBDs on one S-trimer ^17^. Some of the most potent SARS-CoV-2 NAbs target the receptor binding motif (RBM) of the RBD, an area in direct contact with ACE2 ^15^. However, only 8 of the 17 contact residues (47%) in the RBM are conserved between SARS-CoV and SARS-CoV-2 ^18^. Thus, activity of RBM-specific NAbs is likely to be affected by frequently occurring mutations in this region ^19,20^.

In late 2020, SARS-CoV-2 mutational variants of concern (VOC) started emerging that contained mutations in the RBD, which rendered these VOCs (partially) resistant to several earlier obtained neutralizing antibodies ^21^ and decreased the efficiency of current vaccines and therapeutic mAbs ^22–25^. Of particular note was the Beta (B.1.351) VOC, that first showed significant immune escape from serum neutralizing antibodies due to a E484K mutation in the RBD, in combination with N501Y, which was previously described in Alpha ^22,26^. Later Delta (B.1.617.2) displayed increased transmissibility and pathogenicity to an extent where it rapidly became the prevalent circulating isolate ^27^. Since late 2021, we have witnessed a rapid spread of the Omicron (B.1.1.529) VOC, which has since split into several sublineages ^28–30^. The initial Omicron wave was caused by the BA.1 strain, which, compared to the ancestral strain (Wuhan-Hu-1) contains 32 spike mutations: 15 in the RBD of which 9 in the region of direct contact with ACE2 ^31,32^. Around the same time, the BA.2 strain was reported, which quickly took over in many countries, and more recently BA.4 and BA.5, which probably evolved from BA.2, have started to circulate and become globally dominant ^30,33^. BA.2 has additional mutations not present in BA.1, while it lacks others. BA.4 and BA.5 share the same RBD mutations (therefore we address this VOC as BA.4/5), namely L452R (also present in Delta), F486V and wild type amino acid at position Q493 on top of the mutations reported for BA.2 ^34^. The accumulated RBD mutations have greatly diminished the efficacy in preventing infections of available vaccines while most NAbs, including those in clinical use or in development are rendered partly or completely ineffective ^25,33–36^.

One way to evade viral escape is by using a cocktail of antibodies with different specificities to simultaneously target multiple epitopes on the S-trimer. Indeed, in some cases, antibody cocktails where two antibodies showed synergy prevented mutational escape observed with the individual NAbs ^37–40^. Another way to achieve synergy is to create multivalent constructs, such as bispecific antibodies (bsAbs) which contain two different Fab arms that target different spike epitopes. This approach has several advantages. First, bsAbs have shown improved resistance to emerging SARS-CoV-2 variants compared to monospecific NAbs ^41–43^. Second, avidity of a bsAb could give additional benefits by utilizing binding mechanisms often not available to monospecific bivalent mAbs, such as crosslinking two RBDs on one spike (intra-spike) or crosslinking two adjacent spikes on the virion (inter-spike). Thirdly, the development of a single molecule for clinical use might have practical advantages over producing multiple ones for use in a cocktail.

Here, we used a straightforward method to produce several immunoglobulin G (IgG)-like bsAbs from individual NAbs isolated from SARS-CoV-2 Wuhan-Hu-1 infected individuals in early 2020 ^8^. We generated bsAbs by combining potent and highly specific SARS-CoV-2 RBM targeting NAbs COVA2-15 and COVA1-18 with COVA1-16 and COVA2-02, which are less potent but cross-react with SARS-CoV. COVA2-15 can bind both “up” and “down” RBD ^8^ while COVA1-18 was shown to protect against SARS-CoV-2 Wuhan-Hu-1 infection in cynomolgus macaques ^44^. COVA1-16 binds a highly conserved non-RBM epitope but is still able to sterically block ACE2 binding ^45^. COVA2-02 is a less studied NAb targeting a distinct RBD epitope outside of the RBM. Notably, COVA1-16 largely retained its neutralization potency against previously tested SARS-CoV-2 VOCs; while COVA1-18 and COVA2-15 did not neutralize or had significantly (>100-fold) reduced potency against Beta ^22,46^. Overall, our observations on the different binding and neutralization properties of the here generated bsAbs and their monospecific counterparts may contribute to the development of rationally designed antibody-based immunotherapies.

## Results

### Generation of SARS-CoV-2 bsAbs using controlled Fab-arm exchange (cFAE)

Well suited candidates to use as part of multivalent constructs are cross-neutralizing Abs that can neutralize different sarbecoviruses and usually target more conserved areas outside of the RBM. They are relatively rare in comparison with NAbs with narrow specificity, but some of those described have the significant advantage of greater breadth and resistance to viral mutations ^47–49^. As candidates for the generation of our bsAb constructs we chose two such antibodies, COVA1-16 and COVA2-02, in combination with our most potent SARS-CoV-2 NAbs, COVA1-18 and COVA2-15, that are both RBM-binders (Fig 1A). We used controlled Fab arm exchange (cFAE), an efficient method to rapidly produce bsAbs from two parental IgG1 using a redox reaction ^50^. The parental IgG1 molecules contain a single mutation in the Fragment crystallizable (Fc) region of each antibody (either F405L or K409R) to enable heterodimerization and retain correct heavy-light chain pairing after assembly (Fig 1B, ^50,51^). To ensure that F405L and K409R mutations and the cFAE production do not affect Fc effector functions, we performed antibody dependent cellular phagocytosis (ADCP) and antibody dependent cellular trogocytosis (ADCT) assays. The introduced mutations did not adversely affect ADCP and ADCT activity, as we measured similar counts across the panel of antibodies tested, including the monospecific Abs with or without CH3 mutations, and the cFAE bsAbs studied here (Supplemental Fig. 1A,B). Furthermore, introducing either F405L or the K409R mutations did not affect pseudovirus neutralization (Supplemental Fig. 1C). In the remainder of the study, we used antibodies with either F405L or K409R as monospecific Ab controls, and for the sake of brevity, in all figures and text these mutants are labeled only by their COVA names.

**Figure 1.**
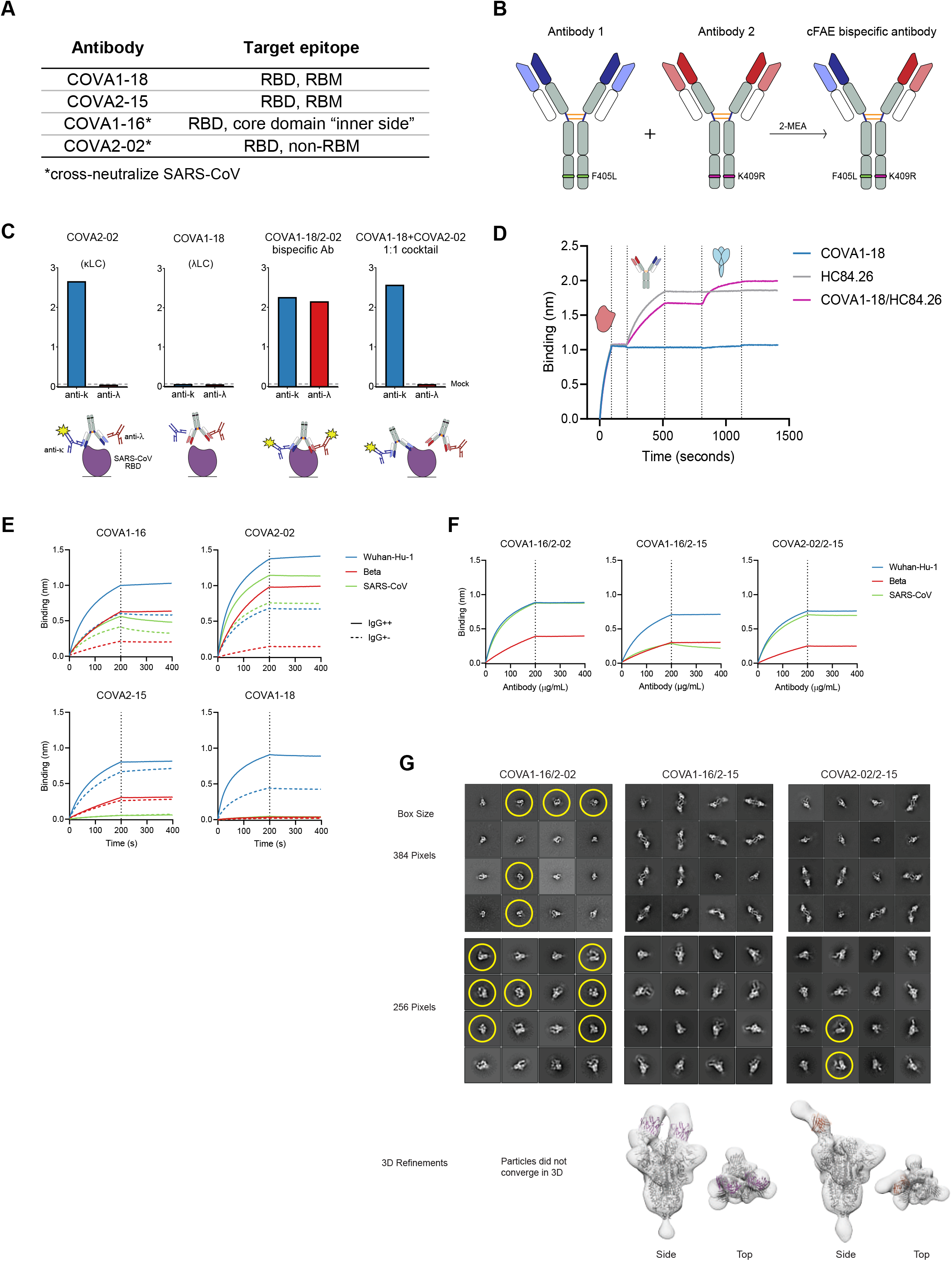
Characterization of IgG1-based bsAbs that bind SARS-COV-2 and SARS-COV S proteins. (**A**) Selected COVA NAbs and their target epitopes. (**B**) Schematic depiction of the method used for bsAb production (cFAE). Matching point mutations (F405L and K409R; EU numbering) are introduced in the parental antibodies, which then undergo Fab arm exchange in the presence of reducing agent 2-MEA, forming IgG-like bsAbs. (**C**) Confirmation of bispecificity by ELISA. His-tagged SARS-CoV RBD was bound to an NiNTA ELISA plate, followed by the bsAbs or mAb controls and secondary Abs specific for either kappa (i.e. COVA-2-02) or lambda LC (i.e. COVA1-18). Detected signals are depicted in the schematic with yellow stars. (**D**) “Sandwich” biolayer interferometry (BLI) traces to confirm concurrent bsAb binding. BsAb or mAb controls were added after his-tagged HCV E2 was loaded onto NiNTA biosensors. Subsequently, SARS-CoV-2 S was added to measure a second association. (**E**) BLI sensorgrams of IgG++ and IgG+- formats of selected COVA NAbs binding to Wuhan-Hu-1, Beta and SARS-CoV S proteins. The dotted lines represent the end of NAb association and the start of dissociation. (**F**) BLI sensorgrams of bsAbs binding to Wuhan-Hu-1, Beta and SARS-CoV S proteins. The curves are representative of two independent experiments. (**G**) Representative 2D class averages from nsEM analysis and corresponding 3D reconstructions, COVA1-16/2-15 and COVA2-02/2-15 bsAbs bound to SARS-CoV-2-6P-Mut7 Spike protein. Due to heterogeneity, particles did not converge to a stable 3D class for COVA1-16/2-02. For the 2D classes, datasets were processed with a box size of 384 pixels to show inter-spike avidity and a box size of 256 pixels to show single Spike proteins. Trimer degradation in the COVA1-16/2-02 and COVA2-02/2-15 complexes are highlighted with yellow circles. Fc portions of the bsAbs in the 2D classes are seen as faint ghost densities near the Fabs. For the 3D refinements, a box size of 256 pixels was used to focus on a single Spike complex. PDB 6VYB (one RBD-up) and a “dummy” (poly alanine) Fv model were fit into the maps.

To test whether cFAE yielded dual-specific antibodies and not a cocktail of monospecific antibodies we used binding enzyme-linked immunosorbent assay (ELISA) and Biolayer interferometry (BLI). We exploited the difference in light chain isotype and binding specificity of COVA2-02 (κ light chain, and binds to Wuhan-Hu-1 and SARS-CoV RBD) and COVA1-18 (λ light chain, only binds to Wuhan-Hu-1 RBD). SARS-CoV RBD was immobilized on the ELISA plate and incubated with the monospecific antibodies, a 1:1 cocktail of COVA2-02 and COVA1-18 or COVA2-02/1-18 bsAb. After detection with secondary antibodies specific for either the κ or λ light chain, only the bsAb showed measurable ELISA signal for both anti-κ and anti-λ, confirming the dual specificity of the bsAb (Fig. 1C). For Octet, we first produced a “dead arm” bsAb by pairing COVA1-18 with HC84.26, an antibody that binds the E2 glycoprotein of the hepatitis C virus ^52^. We then loaded E2 on the BLI sensor, followed by the bsAb, or its parental monospecific controls, and subsequently the Wuhan-Hu-1 S protein. As expected, only the bsAb showed binding to both proteins in this assay (Fig. 1D). These results confirmed that cFAE yielded bonafide bsAbs.

### COVA RBD antibodies rely on avidity for strong binding

Previous studies comparing full IgGs with single Fabs suggest that RBD-targeting NAbs COVA1-16 and COVA1-18 need bivalency for strong binding and neutralization ^17,44,45^. To corroborate these results and determine the influence of bivalency on binding and neutralization potency of all NAb candidates, we produced additional “dead arm” bsAbs by combining COVA-16, COVA2-02 and COVA2-15 with HCV-specific HC84.26. These constructs have the size of an IgG but essentially act as single Fabs, with all potential avidity effects of a fully functional bivalent IgG being eliminated. We compared the binding of the bsAbs containing one irrelevant arm (IgG+-) with their parental counterparts (IgG++) to S proteins of Wuhan-Hu-1, Beta and SARS-CoV by BLI (Fig. 1E). As expected, IgG+- NAbs showed lower binding to Wuhan-Hu-1 S protein than the parental antibodies. This effect was most pronounced for COVA1-16, COVA1-18 and COVA2-02, whereas for COVA2-15 the binding for IgG++ and IgG+- was comparable. This effect was even more pronounced for Beta S, where COVA1-16 and COVA2-02 IgG++ displayed relatively high binding which was substantially decreased for IgG+-, whereas binding of COVA2-15 IgG++ and IgG+- to Beta S was comparable. As previously reported ^46^, COVA1-18 did not bind to Beta S. IgG+- versions of cross-reactive COVA1-16 and COVA2-02 showed slightly lower binding to SARS-CoV S than their IgG++ counterparts, while COVA2-15 and COVA1-18 did not bind SARS-CoV S. These results indicate that COVA NAbs, in particular COVA1-16, COVA2-02 and COVA1-18, need avidity for strong binding to their target.

### BsAbs retain binding to sarbecovirus S proteins

We then performed BLI binding measurements of bsAbs COVA1-16/2-02, COVA1-16/2-15 and COVA2-02/2-15 to SARS-CoV-2 Wuhan-Hu-1, Beta and SARS-CoV S proteins. We did not continue testing COVA1-18, because of it is limited binding breadth (Fig. 1E). All three bsAbs retained binding to the Wuhan-Hu-1, Beta and SARS-CoV S protein (Fig. 1F) and both bsAbs containing COVA2-02 showed substantial binding to SARS-CoV S, despite the COVA2-15 arm not contributing to this interaction.

Next, we used negative stain EM (nsEM) to study the binding mode of the bsAbs to Wuhan-Hu-1 S. We observed intra-spike binding to the RBD for COVA1-16/2-02, COVA1-16/2-15, and COVA2-02/2-15 (Fig. 1G). Interestingly, 2D classes of COVA1-16/2-15 and COVA2-02/2-15 bsAbs showed a wide array of crosslinked poses and antibody stoichiometries, indicating their ability to also bind two spikes simultaneously (inter-spike avidity). Conversely, COVA1-16/2-02 did not display any crosslinking phenotypes. Both COVA1-16/2-02 and COVA2-02/2-15 showed signs of trimer degradation (Fig. 1G). Due to these variabilities, only some particles converged to stable 3D maps. The similarities between COVA1-16/2-15 and COVA2-02/2-15 suggest the COVA2-15 arm is the main contributing factor to the observed avidity effects. All antibodies bound to the RBD, which was confirmed by fitting PDB model 6VYB (1 RBD-up) into the 3D refinements. The 3D refinement of COVA1-16/2-15 showed one fab arm binding to RBD-up and the other to RBD-down, while COVA2-02/2-15 showed a fab binding to RBD-up. However, since these 3D refinements represent only particles that converged in 3D, we presume that the antibodies can bind to both RBD conformations.

### COVA IgG++ and IgG+- NAbs bind the SARS-CoV-2 S protein with diverse stoichiometries

To further assess the binding characteristics of the monospecific IgGs, corresponding “dead arm” bsAbs and the bsAbs we used mass photometry (MP). This single particle mass analysis technique allows for accurate measurements of the highly heterogeneous complexes formed after incubation of antibodies with SARS-CoV-2 Wuhan-Hu-1 soluble S-trimer ^17,53,54^. The measured mass histograms reveal distinct peaks which directly translate to particles representing certain Ab to S binding preferences (e.g. 1:1, 2:1 IgG:S-trimer stoichiometry). Several factors may influence the predominant binding stoichiometry of a particular IgG: the fact that an S-trimer contains three RBD domains, each of which can occupy an “up” or “down” state, the potential of steric hindrance in the binding interaction, and the ability of a bivalent antibody to use avidity in binding, thereby crosslinking multiple domains of the S-trimer simultaneously. As previously shown COVA NAbs have different preferences of binding stoichiometries to the S-trimer (e.g. COVA1-18 has a 1:1 and COVA2-15 a 2:1 binding preference), and these are seemingly uncorrelated to the affinity and neutralization potency of the respective antibodies ^17^. Here we repeated and confirmed the measurements for COVA1-18 and COVA2-15 and determined that IgG++ COVA1-16 binds predominantly with a 1:1 IgG:S-trimer stoichiometry, while COVA2-02 preferentially binds with a 2:1 stoichiometry (Fig. 2A). We observed an increase in free spike for COVA1-16, COVA2-02 and COVA1-18 IgG+- (Fig 2A), but not for COVA2-15 IgG+-, which is in line with the differences observed in biolayer interferometry (BLI) for these antibodies (Fig. 1E). Additionally, for COVA2-02 and COVA2-15 IgG+- we observed an increase in apparent stoichiometry from ∼2 to ∼3 Abs per spike, with a Poisson-like distribution between the different stoichiometries (Fig. 2A). The loss of affinity for IgG+- COVA1-16, COVA2-02 and COVA1-18 compared to IgG++ indicates these mAbs are dependent on intra-spike avidity for successful binding. On the other hand, the lower stoichiometry of IgG++ compared to IgG+- suggests that conventional COVA2-15 uses both arms to bind multiple RBDs in a single spike, meaning it can also utilize intra-spike avidity, but as there is no loss of affinity for COVA2-15 IgG+-, this is not crucial for robust binding. The distribution of particles for each measured IgG bound to the S-trimer, as well as for their “dead arm” counterparts is summarized as a heat map in Figure 2B. We here show that the SARS-CoV-2 NAbs included in this study use intra-spike avidity for efficient binding, i.e. we confirm they utilize both Fab arms to reach their binding potential.

**Figure 2.**
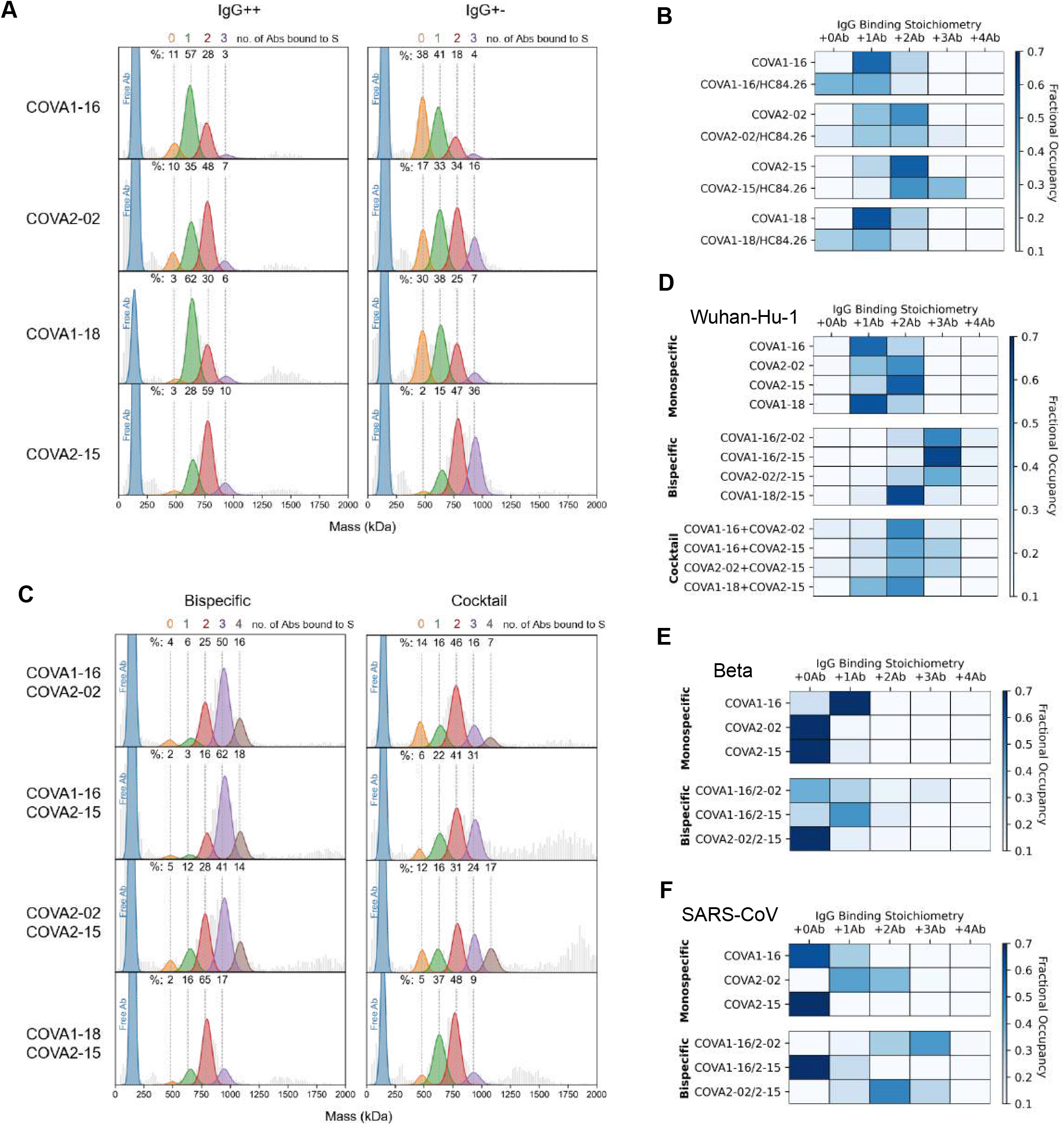
Binding characteristics of bsAbs determined by single particle mass photometry. Mass photometry (MP) measurements of COVA IgG++ and IgG+- formats (**A**), in complex with soluble Wuhan-Hu-1 S protein. The mass histograms were fitted with Gaussian curves corresponding to each distinct assembly. The vertical dashed lines indicate the expected peak positions of each IgG-bound species. All measurements were performed with a 3:1 Ab:S ratio. Percentage values are derived from the normalized summation of two 120 second MP acquisitions. Fractional occupancies of Wuhan-Hu-1 S binding are summarized as heat maps for IgG++ and IgG+- NAbs. (**B**) The darkest blue color for each NAb represents its preferred binding stoichiometry. (**C**) MP measurements of bsAbs and corresponding cocktails to Wuhan-Hu-1 S and the corresponding heatmaps in (**D**). Binding of the bsAbs in comparison to the parental NAbs to SARS-CoV-2 Beta (**E**) and SARS-CoV (**F**) S proteins was assessed and summarized as a table of occupancies.

### BsAbs bind with higher stoichiometries to SARS-CoV-2 and SARS-CoV S proteins

Next, we used mass photometry to determine the binding behavior of the developed bsAbs to the SARS-CoV-2 (Wuhan-Hu-1) S-trimer, in comparison to the parental IgGs and 1:1 cocktails of the parental IgGs. We observed a single peak at the expected size of a human IgG1 (∼150 kDa) for all bsAbs when no S protein added indicating that the cFAE process did not result in antibody aggregation (Supplemental Fig. 2).

When measuring a 1:1 cocktail of monospecific antibodies, the predominant stoichiometry is 2:1 IgG:S-trimer for all mixtures (Fig. 2C, D). However, when measuring COVA1-16/2-02, COVA1-16/2-15 and COVA2-02/2-15 bsAbs, we observed a 3:1 stoichiometry, which was not observed for any of the original COVA mAbs (this study and ^17^) (Fig. 2C, D). A 3:1 binding stoichiometry could be consistent with three bsAbs using one arm each resulting in three of the six potential epitopes being occupied, or with cooperative binding by the three antibodies using both arms, in which case up to six epitopes could be occupied. Another explanation might be that these bsAbs use non-direct avidity or a ligand rebinding mechanism ^55^, meaning they let go with one arm, but quickly latch on again targeting a different epitope. A smaller proportion of the observed complexes (∼15%) represented 4 IgG’s bound to 1 S protein. Again, this could represent 4 antibodies binding with one arm, but may also be consistent with crosslinking two epitopes of the same RBD, something observed before with other SARS-CoV-2 bsAbs ^42^. The stoichiometry did not increase for bsAb COVA1-18/2-15 compared to monospecific COVA2-15 (Fig. 2A), probably because both arms of this antibody target roughly the same epitope (the RBM of the RBD). Interestingly, we observed various distinct particles with masses above 1500 kDa for the COVA1-16/2-15 and COVA2-02/2-15 bsAbs and the corresponding cocktails, which likely represent complexes of two or three S-trimers crosslinked by the antibodies (Supplemental Fig. 3). However, these larger complexes were hardly detected for COVA1-16/2-02 bsAb or the corresponding cocktail. This suggests that the COVA2-15 arm may induce inter-spike binding, corroborating the findings from NS-EM (Fig 1G).

Additionally, we used MP to determine binding stoichiometries of the mAbs and bsAbs to the S-trimer of Beta (Fig 2E) and SARS-CoV (Fig 2F). COVA2-02 and COVA2-15 displayed a stoichiometry close to 0:1 indicating that both antibodies only bind weakly to Beta S, while COVA1-16 retained its binding against this VOC, with a predominantly 1:1 IgG:S-trimer stoichiometry. However, combining COVA1-16 with COVA2-02 or COVA2-15 as bsAbs improved binding to Beta S as we observed an increase to 1:1 stoichiometry for both bsAbs and minor proportions of higher occupancies (2:1 or 3:1) (Fig 2E). As expected, we did not observe binding of COVA2-15 to the SARS-CoV S protein (0:1 stoichiometry). While most of COVA1-16 did not bind SARS-CoV S, ∼30% did bind (1:1) (Fig. 2F). In contrast, the binding of COVA2-02 was heterogeneous, yielding a mix of 1:1 and 2:1 IgG:S-trimer stoichiometries. Combining COVA2-02 with COVA1-16 or COVA2-15 in a bsAb increased SARS-CoV S occupancy to 2 or 3 bsAbs bound per S. This indicates that combining cross-neutralizing antibodies, such as COVA2-02 with other mAbs in a bsAb changes the number of potential binding mechanisms.

### COVA mAbs utilize both arms for potent neutralization

Next, we tested the neutralization activity of the different mAb versions against a panel of pseudoviruses. The mutations present in the RBD of the S protein that could affect the antibody neutralization potency are summarized in Figure 3A.

**Figure 3.**
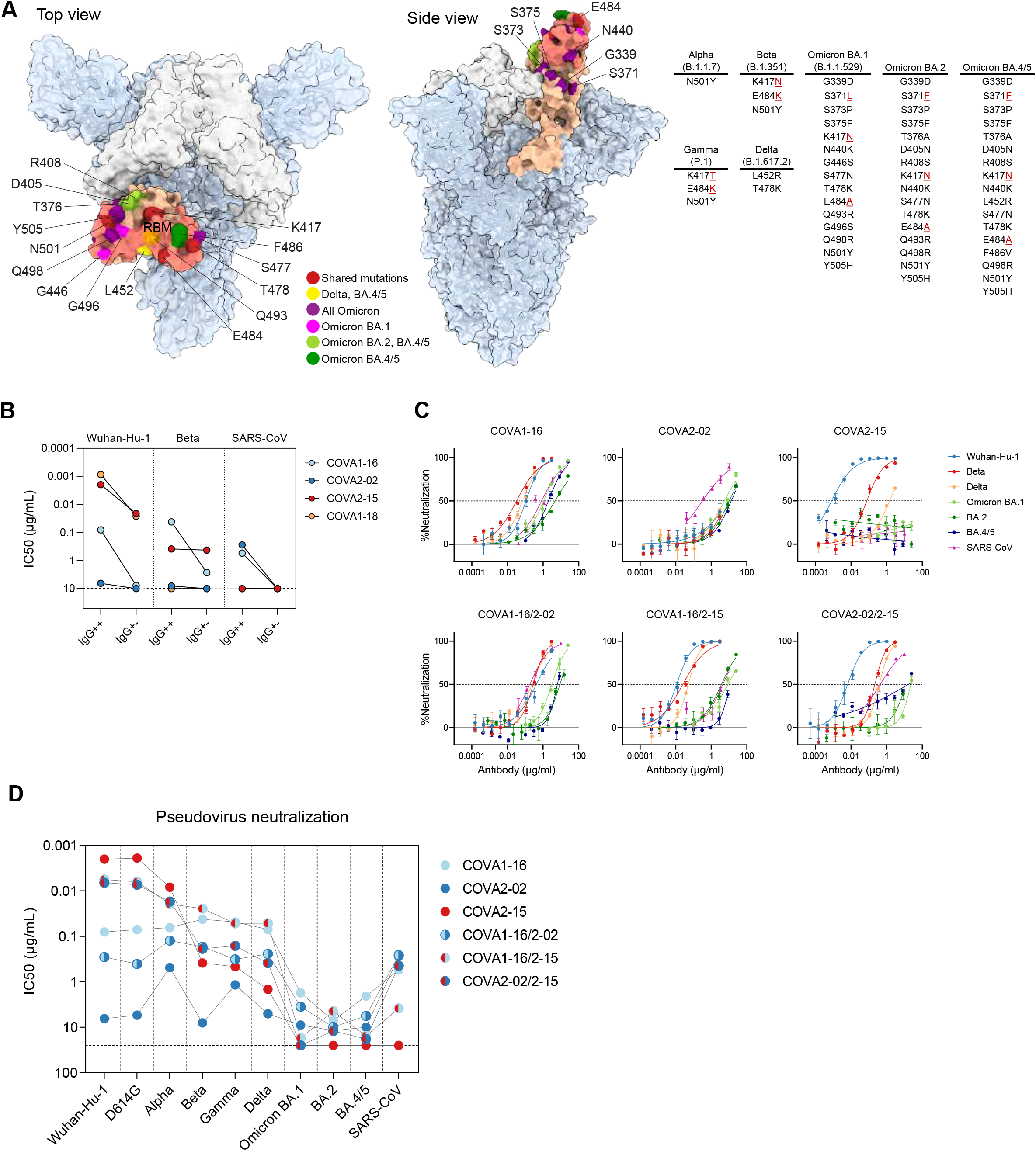
BsAbs potently neutralize SARS-CoV-2 variants and SARS-CoV in pseudovirus neutralization assays. (**A**) Structure of SARS-CoV-2 S trimer with 2 “down” RBD’s (white) and one “up” (salmon) (PDB: 6ZGG). The RBM of the RBD is highlighted in darker salmon. RBD mutations of SARS-CoV-2 variants are listed in a table and pointed out on the RBD structure in red (positions shared between several variants), yellow (mutation shared between Delta and Omicron BA.4/5), purple (mutations shared between all Omicron subvariants), magenta (mutations unique to Omicron BA.1), light green (mutations shared between Omicron BA.2 and BA.4/5) and dark green (mutations unique to Omicron BA.4/5). (**B**) Half maximal inhibitory concentration (IC_50_) values of COVA NAbs (IgG++) or “dead arm” bsAbs (IgG+-) against Wuhan-Hu-1, Beta and SARS-CoV pseudoviruses. In case of IC_50_ >10 μg/mL (indicated by a dotted line) the antibody was considered non-neutralizing. (**C**) Representative neutralization curves of monospecific and bispecific COVA NAbs against SARS-CoV-2 (Wuhan-Hu-1), SARS-CoV-2 variants and SARS-CoV pseudovirus neutralization. The dotted lines indicate 0% and 50% neutralization. Shown is the mean ± SEM of technical triplicates. (**D**) IC_50_ titers of sarbecovirus neutralization by COVA monospecific and bispecific NAbs. In case of IC_50_>25 μg/mL (indicated by a dotted line) the antibody was considered non-neutralizing. Each dot represents the mean IC_50_ value of at least two independent experiments performed in triplicate. Connected dots represent data from the same NAb.

First, we tested if “dead arm” bsAbs lose neutralization potency compared to conventional mAbs. The non-RBM targeting mAbs COVA1-16 and COVA2-02 lost neutralizing activity against Wuhan-Hu-1, Beta and SARS-CoV completely when one arm was paired with HC84.26 (IgG+-) (Fig. 3B). Potent RBM-targeting mAbs COVA2-15 and COVA1-18 with IC_50_’s of 1-2 ng/mL against Wuhan-Hu-1, retained the capacity to neutralize Wuhan-Hu-1 pseudovirus as “dead arm” bsAbs but were 11-fold and 30-fold less potent, respectively. Interestingly, COVA2-15, while being significantly (∼190-fold) weakened by the mutations present in the Beta variant, has the same potency against this VOC as an IgG++ and IgG+- (Fig. 3B, Supplemental Fig. 4A). COVA1-18 did not neutralize SARS-CoV or strains that harbor the E484K mutation including Beta, either as IgG++ or IgG+-. Overall, replacing one of the arms of a COVA IgG with an irrelevant Fab resulted in a significant reduction in neutralization potency.

### BsAbs combine the neutralization potency and breadth of parental antibodies

To evaluate the neutralizing activity of the bsAbs, we compared the breadth and potency to their monospecific counterparts in pseudovirus neutralization assays against SARS-CoV-2 (Wuhan-Hu-1), the main mutational variants (D614G, Alpha, Beta, Gamma, Delta, Omicron BA.1, BA.2 and BA.4/5), as well as SARS-CoV.

Monospecific COVA1-16 showed broad neutralization activity with IC_50_ values of 0.04-0.08 μg/mL against Wuhan-Hu-1 and mutational variants D614G, Alpha, Beta, Gamma and Delta, consistent with earlier findings ^8,45,46^. It was also active against Omicron and its subvariants, albeit with significantly higher IC_50_ values (1.7-7.1 μg/mL) (Fig. 3C, D, Supplemental Fig. 4C). COVA2-02 neutralizes SARS-CoV with similar potency as COVA1-16 (IC_50_ of 0.3-0.6 μg/mL), but only weakly neutralizes SARS-CoV-2 and its variants, including Omicron (IC_50_ values around 10 μg/mL), which is notable considering this NAb was isolated from a SARS-CoV-2 Wuhan-Hu-1 infected individual. On the other hand, COVA2-15 potently neutralized Wuhan-Hu-1, D614G and Alpha, had reduced activity against Beta, Gamma and Delta, but did not neutralize Omicron BA.1, BA.2, BA.4/5 or SARS-CoV. COVA1-18 did not neutralize Beta or SARS-CoV in previous experiments (^36^, Fig 3B, Supplemental Fig. 4A), and was thus not tested against additional SARS-CoV-2 variants.

The COVA1-16/2-02 bsAb was more potent than COVA2-02 alone and overall slightly less potent than COVA1-16 alone (0.5-6-fold) and consistently neutralized all tested viruses with an IC_50_ of 0.1-0.4 μg/mL to most variants, apart from Omicron BA.1 (3.5 μg/mL), BA.2 (9.8 μg/mL) and BA.4/5 (5.6 μg/mL) (Fig. 3C, D, Supplemental Fig. 4C). Combining COVA1-16 with COVA2-15 in a bsAb increased the potency in relation to COVA1-16 alone and broadened the response in relation to COVA2-15 (Fig. 3D, Supplemental Fig. 4C) and potently neutralized most SARS-CoV-2 variants (IC_50_’s of 0.02-0.006 μg/mL) and SARS-CoV (3.8 μg/mL), but exhibited weak neutralization of Omicron BA.1 (17.2 μg/ml) and BA.4/5 (15.5 μg/mL). Interestingly, while not being very potent against Omicron BA.2 (IC_50_ of 4.4 μg/mL), it had slightly better activity against this VOC than each of its components individually (1.6 and 2.7-fold) or a cocktail of the parental antibodies (4-fold). Similarly, we observed that COVA2-02/2-15 bsAb combined the breath of COVA2-02 and the potency of COVA2-15, and neutralized Wuhan-Hu-1, D614G and Alpha efficiently (IC_50_s of 0.02-0.007 μg/mL), was weaker (IC_50_ of 0.2-0.4 μg/mL) against Beta, Gamma and Delta, showed very weak activity against Omicron BA.2 and BA.4/5 (IC_50s_ of 12 and 18 μg/mL, respectively) and no activity against Omicron BA.1. Notably, this bsAb had a higher potency against some variants than either of its parental antibodies. For example, COVA2-02/2-15 neutralized Delta with an IC_50_ of 0.4 μg/mL, compared to 5 μg/mL and 1.5 μg/mL for COVA2-02 and COVA2-15, respectively (Supplemental Fig. 4C). Furthermore, COVA2-02/2-15 also retained neutralizing activity against SARS-CoV, despite having one of its arms replaced by COVA2-15, which is not able to neutralize this virus as a monospecific mAb (Fig. 3C, D, Supplemental Fig. 4C). We also tested cocktails of the two parental monospecific antibodies (1:1 mix) (Supplemental 4B, C). Overall, the cocktails showed similar breadth and potency compared to the bsAbs with some exceptions. For example, COVA2-02/2-15 bsAb was 4-fold more potent than the corresponding COVA2-02+COVA2-15 cocktail against Delta (Fig. 3C, Supplemental Fig 4B, C), while a cocktail of COVA1-16+COVA2-02 was 4-fold more potent than the corresponding COVA1-16/2-02 bsAb against Gamma (Supplemental Fig. 4C). The results were corroborated when tested in neutralization assay using authentic SARS-CoV-2 (Wuhan-Hu-1, Beta and Gamma) (Supplemental Fig. 5A,B), although the neutralization IC_50_ values were higher for all antibodies and antibody combinations compared to the pseudovirus assays (Supplemental Fig. 5C). However, pseudovirus assay results correlated strongly with authentic virus neutralization (Supplemental Fig. 5D). In summary, combining the three monospecific RBD-specific NAbs as bsAbs increases their individual neutralizing potential.

## Discussion

The COVID-19 pandemic has an increased demand for developing NAb formulations to be deployed as therapeutics, especially in immunocompromised patients, and to inform future vaccine design. With the continuing emergence of variants that accumulate mutations that often confer NAb resistance, it is clear that future formulations need to focus on increasing neutralizing breadth to remain effective. For example, some potent mAb (cocktail) therapies such as Casirivimab/Imdevimab (REGN-COV2) or Bamlanivimab/Etesevimab (Eli Lilly) which (partly) target non-conserved epitopes in the RBD have become ineffective against Omicron and its subvariants ^56,57^. A solution to counter immune escape beforehand is to select pan-sarbeco NAbs that target more conserved epitopes such as Sotrovimab/S309, COVA1-16 and COVA2-02 that neutralize SARS-CoV and SARS-CoV-2. These mAbs retained activity against most VOCs, although the potency against Omicron and its subvariants was lower ^58^. In contrast, potent SARS-CoV-2 NAbs COVA1-18 and COVA2-15 did not neutralize SARS-CoV and also showed diminished neutralization against most VOCs, indicating that SARS-CoV neutralization might be a proxy for broader SARS-CoV-2 neutralization.

We describe the generation of several bispecific antibodies which combine the specificities of cross-reacting COVA1-16 and COVA2-02 and highly potent COVA2-15. These bsAbs neutralized all tested SARS-CoV-2 variants, albeit with a lowered potency in some cases, in particular against Omicron. Additionally, both COVA1-16/2-15 and COVA2-02/2-15 retained SARS-CoV neutralization. We demonstrate the potential of multivalent constructs (in this case bsAbs) to combine the breadth, potency and antigenic specificity of their monospecific components, and in some cases exhibit synergistic activity, e.g. COVA2-02/1-15 has a greater neutralization potency against Delta than COVA2-02 and COVA2-15 individually or a 1:1 cocktail of COVA2-02 and COVA2-15.

To gain insight in the binding mechanisms used by these bsAbs, we compared their binding stoichiometries on the S trimer to corresponding 1:1 cocktails of parental mAbs using mass photometry. Interestingly, while bsAbs and cocktails showed very similar neutralization potencies, we observed higher binding stoichiometries for bsAbs compared to cocktails. If assumed that the binding of each Fab is independent, then cocktails should show the same binding behavior as bsAbs, since there is an equal amount of total Fab arms in both bsAb and cocktail preparations. Therefore, the differences in binding stoichiometry between cocktails and bsAbs can be directly attributed to the dependence on connectivity of the two different arms i.e. avidity. We also observed higher stoichiometries for the COVA “dead arm” bsAbs, which act as functional Fabs (Fig 2A,B), but an increased proportion of unbound S protein (indicating decreased affinity) which is not observed for the bsAbs. This suggests that while the bsAbs contain two separate Fabs, intra-spike avidity afforded by having the two Fabs in a single IgG allows for increased affinity and neutralizing potency, which gives bsAbs a potential advantage over monospecific Abs or cocktails.

Additionally, NS-EM analysis suggests COVA1-16/2-15 and COVA2-02/2-15, but not COVA1-16/2-02, are able to simultaneously bind two S proteins and thus (at least partly) rely on inter-spike avidity for binding. Mass photometry measurements also showed that bsAbs having one COVA2-15 arm contain a minor proportion of complexes consisting of two S proteins bound to one or more bsAbs (Supplemental Fig. 3B). Larger proportions of inter-spike connected complexes were observed for the corresponding cocktails, which might be explained by steric clashes of the monospecific components that inhibit intra-spike binding and force one arm to bind another S-trimer. In contrast, for the bsAbs we observe less of these complexes and a decrease in free S compared to cocktails, which might be attributed to increased avidity due to intra-spike binding.

We conclude that by combining Abs such as COVA1-16 and COVA2-02, which need bivalency for strong binding and neutralization, with COVA2-15, that is less affected by removal of one Fab arm and is able to bind both “up” and “down” RBD ^8^, more binding options can be achieved (more stoichiometries and the possibility of both intra and inter-spike avidity).

The method used here to make bsAbs (cFAE) allows for rapid generation of human IgG1-like bsAbs from any antibody pair and is applicable to larger scale manufacturing ^50^, making it a straightforward way to rapidly deploy new bsAb candidates to combat the pandemic. The method could eventually be further simplified, as it was shown that the cFAE reaction can happen by adding reducing agent directly to the supernatant, before protein harvesting, which would eliminate the additional step of purifying 2 antibodies separately before getting the desired bsAb product ^59^

Besides the Fab specificity, it is important for engineered therapeutic IgGs to retain their Fc effector functions for optimal *in vivo* efficacy. We demonstrated comparable levels of ADCP and ADCT for the bsAbs and their parental counterparts, as well as retained neutralization following introduction of Fc mutations necessary for the formation of bsAbs. Our constructs could potentially benefit from additional mutations in the Fc tail, such as LS mutations to increase half-life of the antibodies and improve *in vivo* activity ^60^. It has also been shown that SARS-CoV-2 antibodies with low neutralizing potency but broad binding activity, e.g. COVA2-02, can be Fc engineered to reduce viral spread in live mice ^61^.

In summary, this study provides insights into the generation, binding characteristics and neutralization activity of several potent bispecific SARS-CoV-2 antibodies, which contributes to the development of future bivalent therapeutic candidates. Moreover, we underline the importance of utilizing broad, preferably pan-sarbeco NAbs that target conserved epitopes in the design of multivalent constructs that can withstand viral escape caused by the mutations of new SARS-CoV-2 variants.

## Acknowledgements

We thank Yoann Aldon and Tom Caniels for providing spike proteins and helpful discussions, Sylvie Koekkoek for help with cloning the antibodies, and Wouter Olijhoek and Jacqueline van Rijswijk for help with the pseudovirus neutralization assays. L.R was funded by the AMC as part of the local scientific research incentive policy. J.S is a recipient of a Vidi and Aspasia grant from the Netherlands Organization for Scientific Research (NWO, grant numbers 91719372 and 015.015.042). R.W.S is a recipient of a Vici grant from the Netherlands Organization for Scientific Research (NWO, grant number 91818627). This research was supported by an Amsterdam institute for Infection and Immunity Postdoctoral grant (K.S.). A.J.R.H. acknowledges funding by the Netherlands Organization for Scientific Research (NWO) through the Spinoza Award SPI.2017.028. This work was supported by the Fondation Dormeur, Vaduz (R.W.S and (M.J.v.G.).

## Author contributions

Conceptualization, K.S., V.Y., M.J.v.G., R.W.S., J.S.; Methodology, L.R., K.S., V.Y., M.B., A.I.S., J.L.T., J.A.B., M.P.; Investigation, L.R., V.Y., M.B., J.C., A.I.S., J.L.T., S.B., J.A.B., M.P., I.B., J.H.B., D.G.; Resources, L.R., K.S., M.B., J.C., M.J.v.G.; Data curation, L.R., V.Y.; Writing-Original draft: L.R., K.S.; Writing-Review and editing: all authors; Visualization: L.R., V.Y.; Supervision: K.S., D.E., A.B.W., A.J.R.H., M.J.v.G., R.W.S., J.S.; Funding acquisition: A.B.W., A.J.R.H., M.J.v.G., R.W.S., J.S.

## Supplemental figures

**Supplemental Figure 1.**
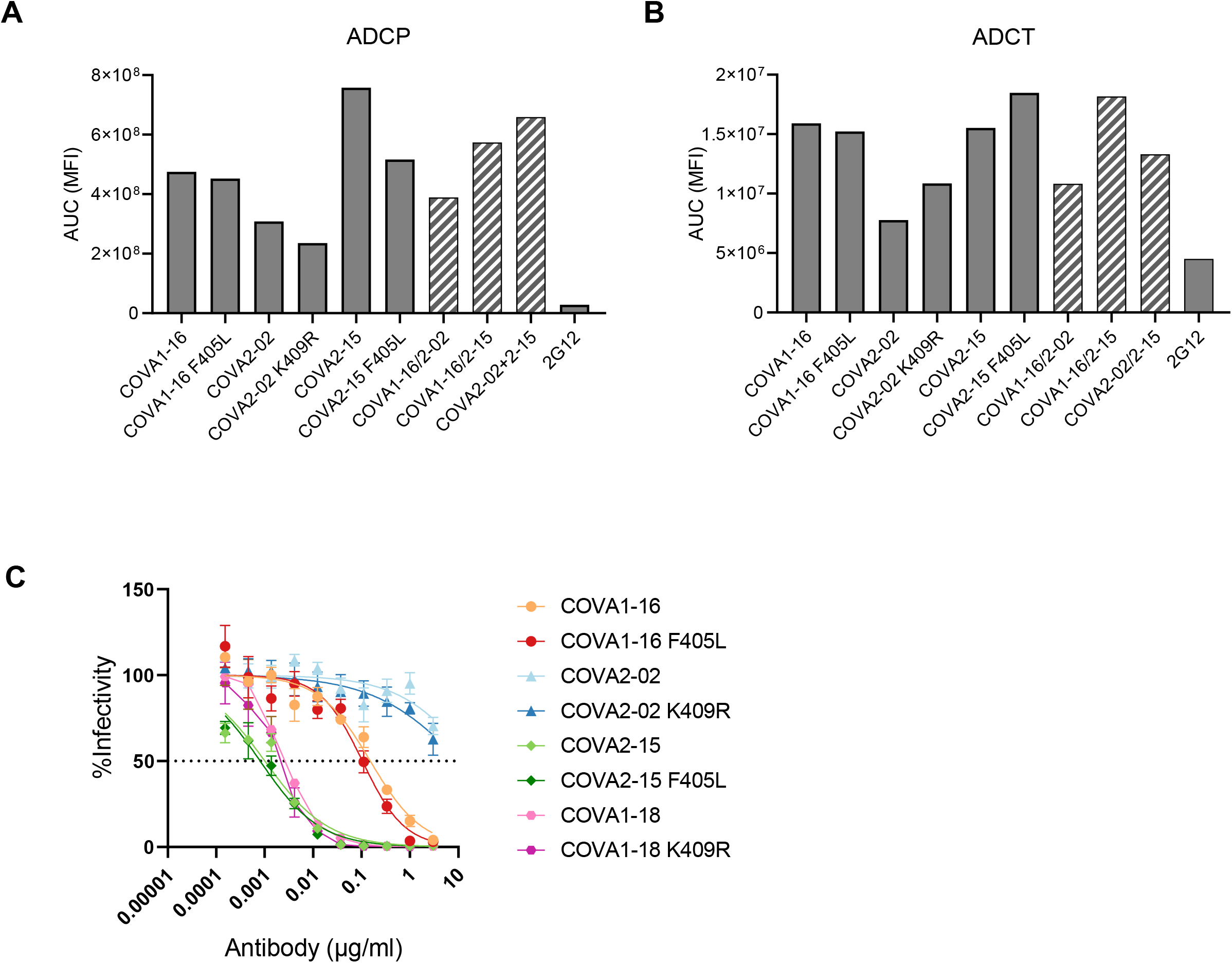
Fc effector functions of bsAbs are retained. Antibody-dependent cellular phagocytosis (ADCP) (**A**) and antibody dependent cellular trogocytosis (ADCT) (**B**) assays were performed and signal was measured using flow cytometry. 2G12, an HIV-1 gp120 specific IgG was used as negative control. Bars represent the area under the curve (AUC) values of the mean fluorescence intensity (MFI). (**C**) SARS-CoV-2 (Wuhan-Hu-1) pseudovirus neutralization curves of COVA monospecific NAbs and versions containing the F405L and K409R mutations needed for bsAb generation by cFAE. The dotted line indicates 50% infectivity. Curves are representative of two separate experiments performed in triplicate.

**Supplemental Figure 2.**
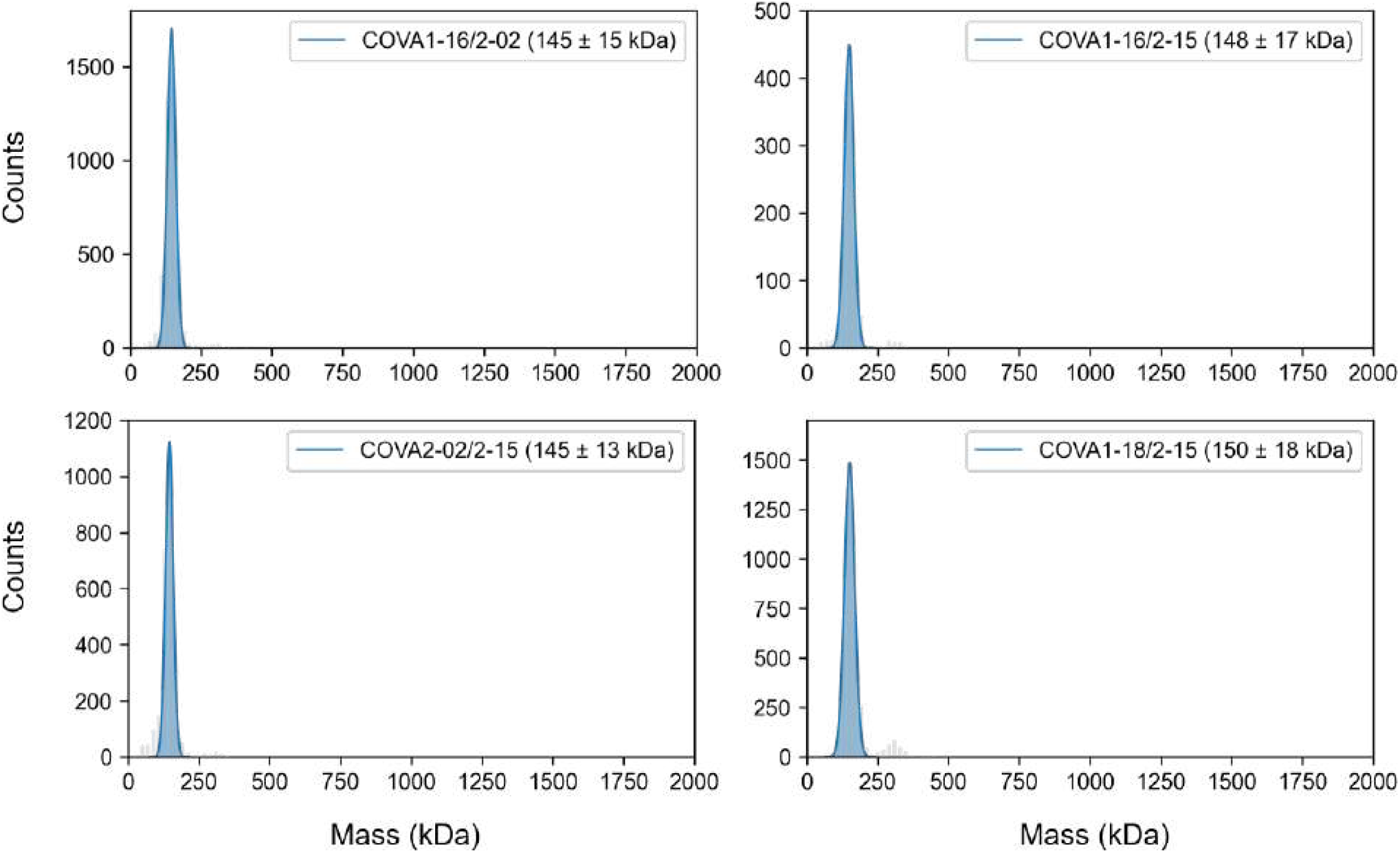
Mass photometry histograms of COVA bsAbs. A single mass distribution at ∼150 kDa is observed for all bsAbs, corresponding to the predicted masses of full IgG1s, confirming that the Fab-arm exchange induced formation of bsAbs does not lead to Ab aggregation.

**Supplemental Figure 3.**
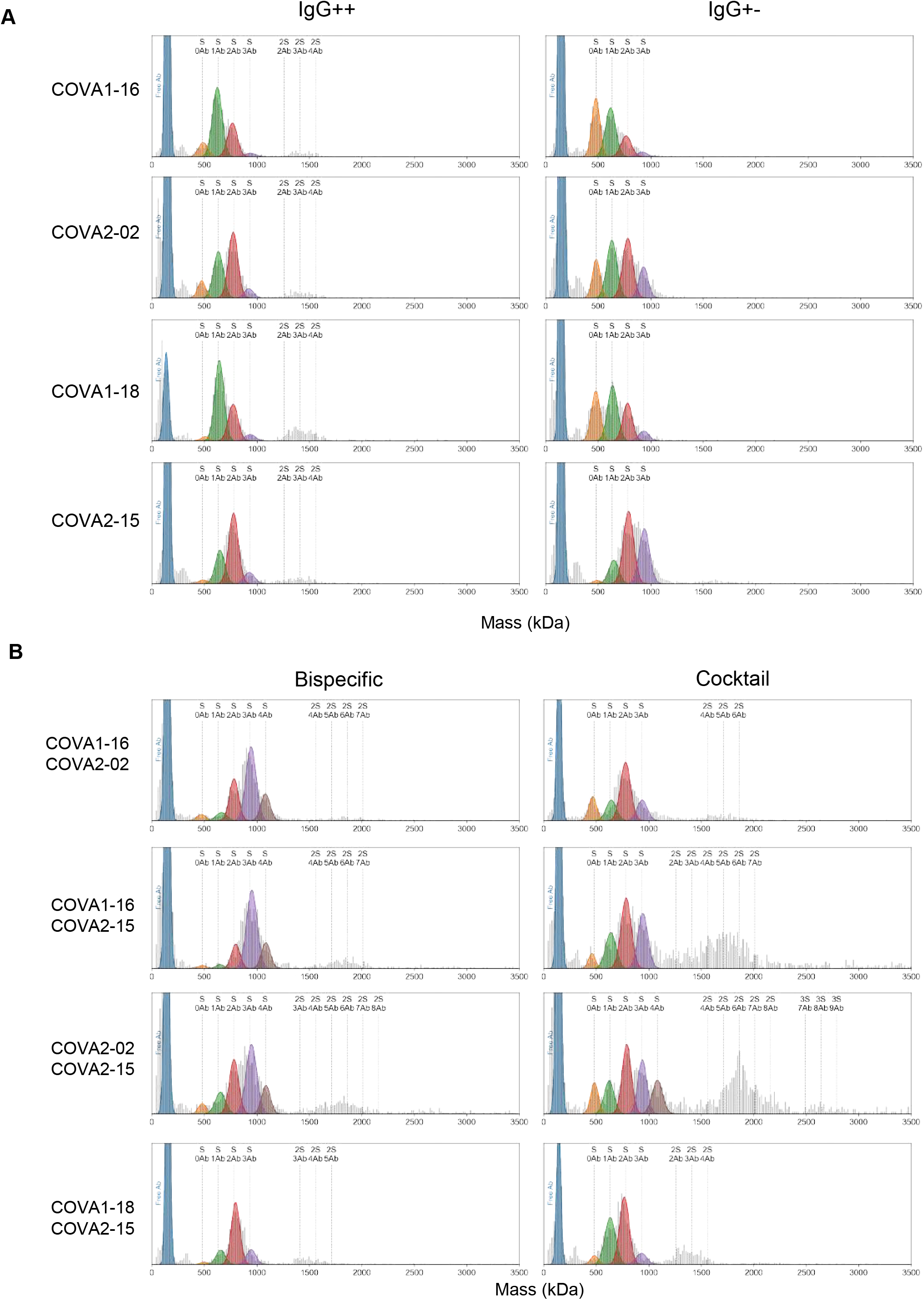
Full MP histograms of COVA monospecific, bispecific and cocktail binding to Wuhan-Hu-1 S. Raw histograms are shown together with the fitted curves for COVA IgG++ and IgG+- constructs (**A**) and COVA bsAbs and cocktails (**B**). Besides the complexes of 1-3 Abs to one S trimer, higher stoichiometries such as 2S:2Ab, 2S:3Ab are indicated with dotted vertical lines where applicable.

**Supplemental Figure 4.**
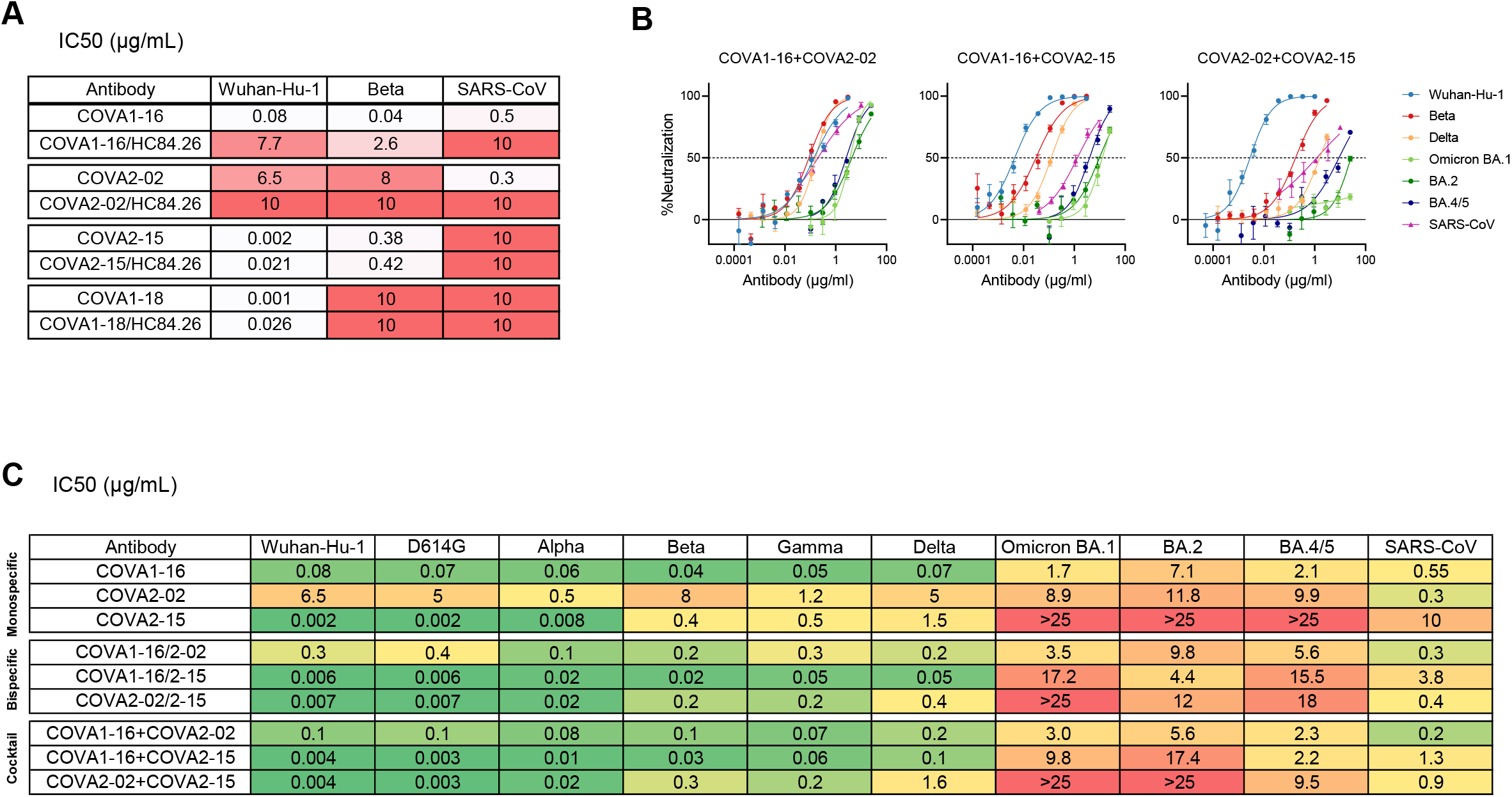
Pseudovirus neutralization of SARS-CoV-2 variants and SARS-CoV by COVA monospecific antibodies, bispecific antibodies and cocktails. (**A**) IC_50_ values (μg/mL) of COVA monoclonal IgGs in comparison to respective “dead arm” bispecifics. All IC_50_S greater than 10 μg/mL were rounded up to 10 and were considered non-neutralizing. (**B**) Representative neutralization curves of 1:1 cocktails of COVA NAbs against SARS-CoV-2 (Wuhan-Hu-1), SARS-CoV-2 variants and SARS-CoV. The dotted lines indicate 0% and 50% neutralization. Data points represent the mean ± SEM of technical triplicates. (**C**) Summary of all IC_50_ values of sarbecovirus neutralization by COVA monospecific and bispecific NAbs and corresponding cocktails. Every value represents the mean IC_50_ of at least two independent experiments performed in triplicate.

**Supplemental Figure 5.**
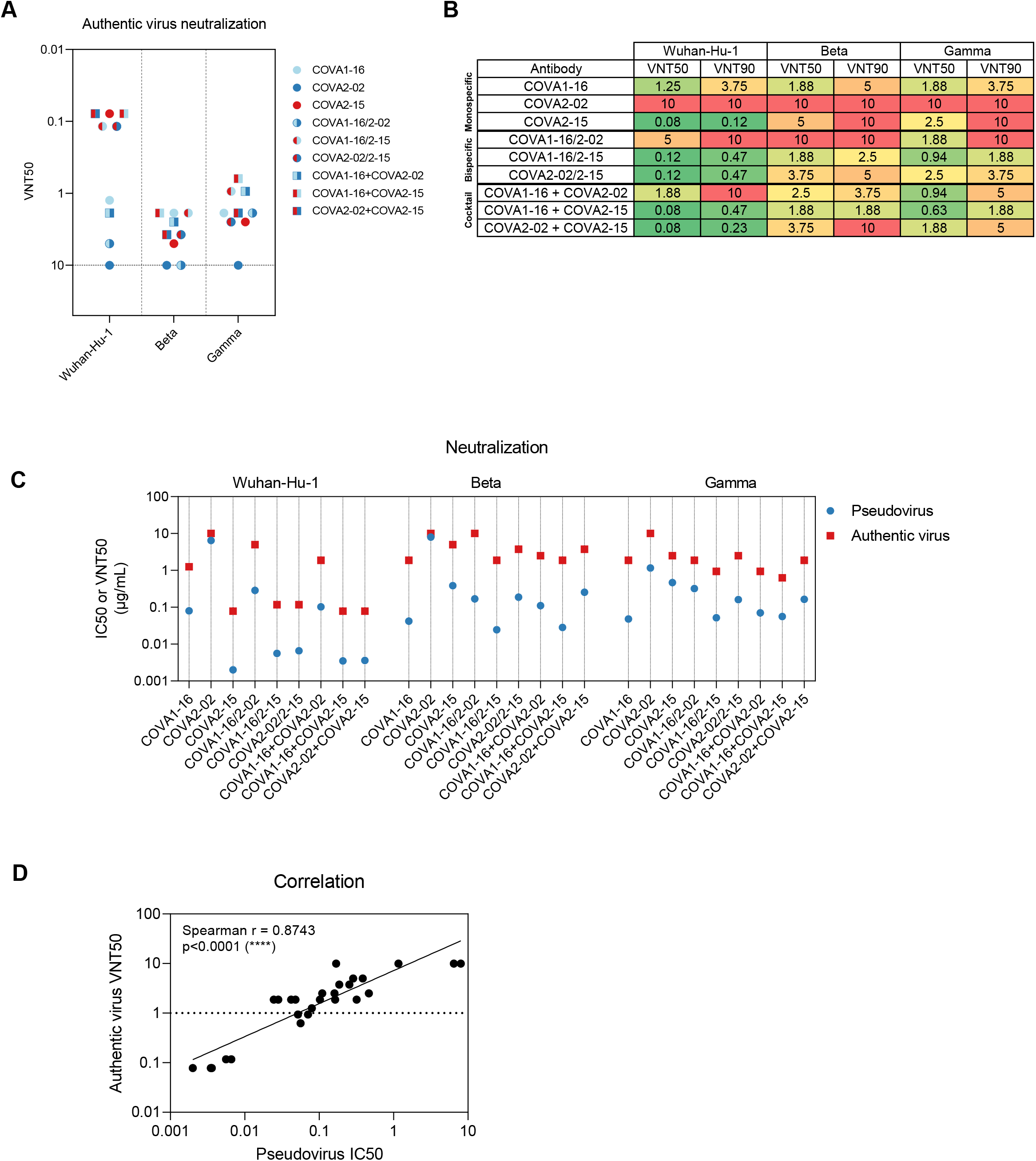
Wuhan-Hu-1, Beta and Gamma authentic virus neutralization. (**A**) Half-maximal virus neutralization titers (VNT_50_) of authentic virus neutralization by COVA monospecific (dots of one color) and bispecific (dots of two colors) antibodies and corresponding cocktails of parental NAbs (squares of two colors). Each symbol represents the mean VNT_50_ value of two replicate measurements. (**B**) Table with summarized VNT_50_ and 90% virus neutralization titers (VNT_90_) of Wuhan-Hu-1, Beta and Gamma neutralization. (**C**) Comparison of IC_50_ (pseudovirus neutralization) and VNT_50_ values (authentic virus neutralization). Each dot represents the mean value from at least 2 separate experiments (IC_50_ values) or 1 experiment performed in duplo (VNT_50_ values). (**D**) Correlation plot of pseudovirus and authentic virus neutralization titers compared for different antibodies and viral strains. Spearman correlation (r) and p value were determined in GraphPad Prism 8.3.0.

## Methods and materials

### Constructs

Prefusion SARS-CoV-2 S was made as described before ^8^. In brief, a gene encoding residues 1-1138 (Wuhan-Hu-1, Genbank: MN908947.3) with proline substitutions at positions 986 and 987 and a GGGG substitution at positions 682-685 was cloned into a pPPI4 plasmid backbone containing a T4 trimerization domain follow by a hexahistidine tag. Prefusion SARS-CoV-2 Beta was produced by introducing the appropriate mutations in the above described construct, as described in ^46^. Prefusion SARS-CoV S was produced as described before ^62^ and was cloned into the same pPPI4 backbone as described above. SARS-CoV-2-6P-Mut7 for nsEM was made as described before ^63^. Briefly, the mutagenesis was performed on the SARS-CoV-2-6P plasmid to include Mut7 (V705C and T883C). The receptor-binding domain (RBD) (residues 319-541) of the SARS-CoV-2 S protein (GenBank: QHD43416.1) and the RBD (residues 306-527) of the SARS-CoV S protein (GenBank: ABF65836.1) were produced as previously described ^8^. The codon-optimized sequence of His-tagged E2 (Genscript Biotech) based on an HCV variant of the genotype 1a H77 strain (Genbank: ABN11232.1 (amino acids 384–659) and GenBank: AF009606.1 (amino acids 660–715)) was cloned into a mammalian expression plasmid. The COVA NAbs used in this study were isolated from participants in the COSCA study, and the variable V(D)J-regions of the heavy and light chain of the antibodies were cloned into corresponding expression vectors containing the constant regions of the human IgG1 as previously described ^8^. The variable heavy and light chain sequences of HC84.26 ^52^ were codon optimized and cloned into a mammalian expression vector as described before ^64–66^ (Genscript Biotech). A Quickchange site-directed mutagenesis kit (New England Biolabs) was used to introduce the F405L and K409R mutations in the heavy chain plasmids of the IgGs.

### Expression and purification of viral proteins and monospecific antibodies

Viral proteins were produced in HEK293F suspension cells (ThermoFisher) and purified as previously described ^8^. Recombinant SARS-CoV-2-6P-Mut7 S protein for nsEM was produced and purified as previously described ^63^. His-tagged E2 was purified by affinity purification using Ni-NTA agarose beads followed by size-exclusion chromatography. Monoclonal antibodies were produced as previously described ^8^.Briefly, after co-transfection of the HC and LC plasmids in a 1:1 ratio and harvest after 5 days, the filtered supernatant was run over 10 mL protein G columns (Pierce). After elution, the purified antibodies were buffer exchanged to PBS using 100kDa Vivaspin6 columns. The IgG concentration was measured on a NanoDrop One (Thermofisher), diluted with PBS to 1mg/mL and stored at 4°C before performing the cFAE protocol.

### Generation of bsAbs by controlled Fab-arm exchange (cFAE)

Equimolar amounts of IgG1-F405L and IgG1-K409R antibodies (1 mg/mL) were mixed with freshly prepared 2-mercaptoethylamine (2-MEA; Sigma) solution (750 mM, pH 7.4), with a volume of 10% of the total reaction volume on a rotating laboratory mixer at room temperature. The mixture was incubated for 5h at 31°C on a thermoblock (Eppendorf), followed by the removal of 2-MEA by buffer-exchanging to PBS using 100kDa Vivaspin6 columns (Sartorius) and storing overnight at 4 °C to allow for reoxidation of the disulfide bonds.

### Enzyme-linked immunosorbent assay (ELISA) for determining bispecificity of bsAbs

His-tagged RBD of SARS-CoV S was diluted to a concentration of 0.8 μg/mL in Tris-buffered saline (TBS) and immobilized on NiNTA 96-well plates (Qiagen) for 2h at RT. Next, monospecific, bispecific IgGs or a 1:1 mix of parental monospecific antibodies were added to the wells, with a concentration of 2 μg/mL in casein, and incubation was allowed for 1.5h at RT. Then, 1:10000 dilutions in casein of either anti-kappa LC or anti-lambda LC secondary antibodies (Bethyl Laboratories) were added for 45 min at RT. In between all abovementioned steps, the plates were washed three times with TBS. After the last step the plates were washed five times with TBS/0.05% Tween-20 and developed with a solution of 1% 3,3’,5,5’-tetramethylbenzidine (Sigma-Aldrich), 0.01% hydrogen peroxide, 100 mM sodium acetate and 100 mM citric acid for 5 min, before adding 0.8 M sulfuric acid to end the development reaction. Optical density (OD) at 450 nm was measured using a spectrophotometer (BMG Labtech).

### Biolayer interferometry (BLI)

All BLI experiments were performed using an Octet K2 instrument (ForteBio). To confirm the bispecificity of the bsAbs we first loaded monomeric his-tagged E2 (10 μg/mL) on Nickel– nitrilotriacetic acid (NiNTA) biosensors (ForteBio) with a binding threshold of 1nm, followed by a wash (120 s) with running buffer (PBS, 0.002% Tween, 0.01% bovine serum albumin) to remove excess protein. Next, the biosensors were dipped into a well with 5 μg/mL bispecific or monospecific NAb for 200 s to measure association, followed by immersion in running buffer to allow dissociation for another 200 s. Finally, the biosensors were moved to a well with Wuhan-Hu-1 S (5 μg/mL) for 200 s, and subsequently to a well with running buffer for 200 s to measure dissociation. To measure binding strength of monospecific and bispecific antibodies, protein A sensors (ForteBio) were first loaded with 10 μg/mL of NAb in running buffer until a threshold of 1nm was reached. After a wash (20 s) in running buffer, the biosensors were submerged in a well with Wuhan-Hu-1, Beta or SARS-CoV S (20 μg/mL) in running buffer for 200 s to measure association, followed by immersion in a well with running buffer for 200 s to measure dissociation of the S-NAb complexes.

### Negative stain-EM sample preparation, data collection, and processing

For each antibody, 0.5 M excess of bispecific IgG was added to stabilized prefusion SARS-CoV-2-6P-Mut7 S protein 30 minutes prior to deposition onto carbon-coated 400-mesh copper grids. The grids were stained with 2% (w/v) uranyl-formate for 90 seconds. Grids were imaged on a Tecnai T12 Spirit at 120 KeV using a 4kx4k Eagle CCD camera. Micrographs were collected using Leginon and the images were transferred to Appion for processing ^67,69^. Particles were picked using a difference-of-Gaussians picker (DoG-picker) ^71^. Data was processed in RELION 3.0 for 2D and 3D classification and 3D refinements ^73^. Figures were generated using UCSF Chimera ^75^.

### Pseudovirus production

HEK293T cells grown in Dulbecco’s modified Eagle’s (DMEM) medium (Gibco) and supplemented with 10% fetal bovine serum, penicillin (100 U/ml) and streptomycin (100 μg/ml) were transfected with pHIV-1_NL43_ΔENV-NanoLuc reporter virus plasmid and a plasmid containing the appropriate S protein. SARS-CoV-2 Wuhan-Hu-1 and D614G, and SARS-CoV S plasmids were made as described in ^64^; SARS-CoV-2 Alpha, Beta, Gamma, Delta and Omicron BA.1, BA.2 and BA.4/5 S plasmids were made as described in Caniels et al. 2021. Supernatant containing the pseudovirus was harvested 48h after transfection. Supernatant was centrifuged at 500 x *g* for 5 minutes and sterile filtered through a 0.22 μm PVDF syringe filter. Pseudovirus was stored at -80°C.

### Pseudovirus neutralization assay

Neutralization assay was performed as described before ^64^. In brief, HEK293T/ACE2 cells were seeded in 96-well culture plates. After 24h, Abs were serial diluted in cell culture medium (DMEM supplemented with 10% fetal bovine serum, penicillin (100 U/ml), streptomycin (100 μg/ml) and GlutaMAX (Gibco)) and mixed in a 1:1 ratio with pseudovirus, and incubated at 37°C for 1 hour. Next, Ab and pseudovirus mixes were added to the cells and incubated for 48 hours. Afterwards, cells were lysed and luciferase activity was measured in the lysates.

### Authentic virus neutralization test

We tested mAbs and bsAbs for their neutralization capacity against the ancestral SARS-CoV-2 virus (German isolate; GISAID ID EPI_ISL 406862; European Virus Archive Global #026V-03883) and VOCs, as previously described ^68^. Briefly, samples were serially diluted in Dulbecco modified Eagle medium supplemented with NaHCO3, HEPES buffer, penicillin, streptomycin, and 1% fetal bovine serum, starting at a dilution of 10 μg/mL in 50 μl.

Subsequently, 50 μL of virus suspension were added to each well and incubated at 35°C for 1 h. Vero E6 cells were added in a concentration of 20,000 cells per well and subsequently incubated for 48 hours at 35°C. After incubation, cells were fixed with 4% formaldehyde/phosphate-buffered saline (PBS) and stained with a nucleocapsid targeting monoclonal antibody. Bound Ab as a measure for infected cells was detected using horseradish peroxidase–conjugated goat anti-human IgG (1:3000) in 2% milk/PBS for 1 hour at RT. After washing, the color reaction was developed using 3,3′,5,5′-tetramethylbenzidine substrate (Thermo Scientific Scientific). The reaction was stopped by adding 0.8 N sulfuric acid, and OD450 (optical density at 450 nm) was measured using standard equipment.

### Mass Photometry

MP experiments were performed using a Refeyn OneMP mass photometer (Refeyn Ltd.). Measurements were mass-calibrated using an in-house prepared protein standard mixture: IgG4Δhinge-L368A (73 kDa ^70^, IgG1-Campath (149 kDa), apoferritin (479 kDa), and GroEL (800 kDa). MP data were processed using DiscoverMP (Refeyn Ltd.). Peaks for each mass species were manually identified and fitted using SciPy ^72^. All MP histograms were plotted using 20 kDa bin widths. Spike/NAb binding experiments were performed as previously described ^17^. In brief, measurements were performed in Tris buffer (25 mM Tris, 100 mM NaCl, pH 7.6 (Sigma-Aldrich)). For each experiment, a 100 nM solution of soluble S protein was mixed with an equal volume of ligand to the desired concentration ratio and incubated at room temperature (22 °C) for 5 min. Afterward, 3 μL of the reaction mixture was immediately transferred to the instrument for measurement.

### Antibody-dependent cellular trogocytosis (ADCT)

HEK-293T cells (Invitrogen) were transfected using SARS-CoV-2 S expression plasmid expression vector and lipofectamine (thermofisher) in OptiMEM as previously described ^74^. SARS-CoV-2 S expression HEK293T cells were stained with PKH26 dye according to the manufactures protocol (Sigma-Aldrich). Next, the stained HEK293T cells were incubated for 30 min at 37°C with serial antibodies dilutions. 2G12-IgG1, specific for HIV-1 gp120, was used as a negative control. After incubation, the cells were washed and THP-1 cells (ATCC), stained with carboxyfluorescein succinimidyl ester according to manufactures protocol (Thermofisher), were added to the HEK293T cells at a 2:1 effector:target cell ratio. The plates were spun down for 30 sec to promote cell to cell contact before incubation for 1 hours at 37°C. After incubation, the plates were washed twice, resuspended in PBS 2% fetal calf serum (FCS) and analyzed using flow cytometry. Trogocytic activity was calculated by the MFI of the double positive PKH26+ CFSE+, THP-1 cells and depicted as the area under the curve.

### Antibody-dependent cellular phagocytosis (ADCP)

The ADCP assay was performed as described previously ^76^. In short, Fluorescent Neutravidin beads (Invitrogen) were coated with biotinylated SARS-CoV-2 S or RBD protein overnight at 4 °C. Next, the beads were washed twice using PBS 2% bovine serum albumin (BSA) and resuspended in PBS 2% BSA at a 1:500 dilution. Serial antibodies dilutions were incubated for 2 hours at 37 °C with 50 μl of the coated beads in a V-bottom 96-well plate. 2G12-IgG1, specific for HIV-1 gp120, was used as a negative control. After incubation, plates were washed and 5×10^4^ THP-1 effector cells (ATCC) were added to each well. The plates were spun down for 30 sec to promote beads to cell contact before incubation for 5 hours at 37°C. Afterwards, the plates were washed twice, resuspended in PBS 2% FCS and analyzed by flow cytometry. The phagocytic activity was determined by the area under curve of the MFI (beads positive cells x mean MFI FITC).

### Statistical analysis and data visualization

All midpoint mAb inhibition concentrations (IC_50_ values) were determined, and data visualization and statistical analyses were performed in GraphPad Prism 8.3.0. Model of the SARS-CoV-2 S protein with VOC mutations was visualized using UCSF ChimeraX ^77^.

